# Integrative modeling of read depth and B-allele frequency improves single-cell copy number calling from targeted DNA sequencing panels

**DOI:** 10.64898/2026.03.12.711292

**Authors:** Dong Pei, Rachel Griffard-Smith, Brahian Cano Urrego, Emily Schueddig

## Abstract

Copy number variations (CNVs) drive cancer initiation and progression, but resolving them at single-cell resolution from targeted DNA sequencing panels remains challenging. The Mission Bio Tapestri platform generates two complementary signals for CNV inference: sequencing depth and B-allele frequency (BAF) from heterozygous variants; however, existing methods such as karyotapR rely primarily on read depth, potentially missing allele-specific events invisible to depth-only approaches. Here we introduce scPloidyR, a hidden Markov model (HMM) that jointly models read depth and BAF at amplicon resolution for single-cell copy number calling from Tapestri data. scPloidyR fits independent per-chromosome Markov chains with copy number states as hidden variables, factorizes emission probabilities into depth and BAF likelihoods, and learns parameters via Baum–Welch expectation-maximization with Viterbi decoding. We compared scPloidyR with the established karyotapR Gaussian Mixture Model (GMM) through two simulation studies that evaluates BAF noise, variant density, amplicon density, sample size, and heterozygosity rate, and through application to a public Tapestri five-cell-line mixture dataset. In simulations, scPloidyR substantially outperformed karyotapR on class-balanced metrics (macro-F1: 0.472 vs. 0.264; alteration F1: 0.902 vs. 0.383 in simulation study 1) when allelic information was available. Adding just one heterozygous variant per amplicon increased scPloidyR accuracy from 0.548 to 0.899 for copy number gains. However, when BAF information was absent, karyotapR outperformed scPloidyR, and high BAF noise substantially degraded joint-model performance. On real data, scPloidyR produced more spatially coherent and biologically plausible copy number profiles. These results establish that joint depth-BAF modeling provides a clear advantage for single-cell CNV calling when allelic information is available, while depth-only methods remain preferable when such information is absent.

**Author Summary:** Cancer cells frequently gain or lose copies of DNA segments, and detecting these changes in individual cells is critical for understanding how tumors evolve and resist treatment. A technology called Tapestri sequences DNA from thousands of single cells and produces two types of signals: how much DNA is present (read depth) and which version of each gene a cell carries (allele information). Existing tools mainly use the first signal, potentially missing important changes that only the second signal can reveal. We developed scPloidyR, a statistical method that combines both signals to more accurately identify DNA copy number changes in single cells. Through simulations and analysis of real cancer cell data, we found that using both signals together substantially improves detection when allele information is available — even a small amount of allele data makes a meaningful difference. However, when allele information is absent, the simpler depth-only approach performs better. Our work provides researchers with a new tool and practical guidance on when each approach is most effective for studying cancer at single-cell resolution.

## Introduction

Copy number variations (CNVs), which include genomic gains or losses ranging from kilobases to entire chromosomes, are frequent drivers of cancer initiation and progression (Zack et al., 2013). Genome-wide aneuploidy, gains or losses of whole chromosomes, is present in approximately 90% of solid tumors and 50–70% of hematopoietic cancers, underscoring its pervasive role in tumorigenesis (Ben-David & Amon, 2020; Taylor et al., 2018). Somatic CNVs activate oncogenes through amplification and inactivate tumor suppressors through deletion, and recurrent patterns of copy number alteration have been cataloged across dozens of cancer types (Zack et al., 2013; Steele et al., 2022). Despite their clinical importance, bulk sequencing averages CNV signals across millions of cells, masking the intratumor heterogeneity that drives treatment resistance and clonal evolution (Navin et al., 2011). Single-cell DNA sequencing overcomes this limitation by resolving copy number profiles at the level of individual cells, enabling reconstruction of clonal architectures and identification of rare subpopulations (Navin et al., 2011; Mallory et al., 2020).

The Mission Bio Tapestri platform is a droplet-based microfluidic system that performs targeted single-cell DNA sequencing of thousands of cells per run (Pellegrino et al., 2018). By amplifying a defined panel of genomic loci, Tapestri generates two complementary signals for CNV inference: sequencing depth (amplicon read counts), which reports total DNA content, and B-allele frequency (BAF) from heterozygous single-nucleotide polymorphisms, which provides allelic information that can distinguish copy number states with similar total counts (Pellegrino et al., 2018). The platform has been widely adopted for profiling hematologic malignancies and solid tumors at single-cell resolution (Pellegrino et al., 2018; Miles et al., 2020; Morita et al., 2020; Albertí-Servera et al., 2021; Zhang et al., 2023).

Several computational tools have been developed for analyzing Tapestri data. The optima R package provides an integrated toolkit for processing Tapestri single-cell multi-omics data, including data import, normalization, quality filtering, and visualization (Pei et al., 2023). karyotapR extends this ecosystem with a GMM approach to copy number calling: it normalizes amplicon read counts against a set of reference cells, simulates synthetic cells at each copy number state using Weibull distributions fitted to reference data, and classifies cells by posterior probability under Gaussian components (Mays et al., 2025; Mays, n.d.). karyotapR has been shown to detect chromosome- and arm-scale aneuploidy across thousands of single cells (Mays et al., 2025). However, karyotapR classifies each amplicon independently without exploiting the spatial ordering of loci along chromosomes, and it relies primarily on read depth, leaving BAF information underutilized and potentially missing allele-specific events such as copy-neutral loss of heterozygosity.

Hidden Markov models (HMMs) provide an alternative framework for copy number inference because they exploit the spatial ordering of genomic loci along chromosomes (Rabiner, 2002). HMM-based methods model copy number states as hidden variables that transition rarely between adjacent loci, with observed signals (depth and/or BAF) emitted according to state-dependent distributions. PennCNV demonstrated this approach on SNP arrays by jointly modeling log R ratio and B-allele frequency within a six-state HMM (Wang et al., 2007), and HoneyBADGER extended HMM-based CNV detection to single-cell RNA-seq by integrating allelic and expression signals through a Bayesian hierarchical framework (Fan et al., 2018). These methods have been successfully applied to array CGH (Shah et al., 2006), SNP genotyping arrays (Wang et al., 2007), and single-cell transcriptomic data (Fan et al., 2018), but HMM-based approaches have not been systematically adapted for targeted single-cell DNA platforms like Tapestri, where amplicon panels are sparse and unevenly distributed.

Integrating BAF alongside read depth can improve CNV detection by resolving allele-specific events, such as copy-neutral loss of heterozygosity and unbalanced gains, that are invisible to depth-only methods (Zaccaria & Raphael, 2021; Wang et al., 2007). However, the value of BAF information depends on the availability of heterozygous variants within the targeted panel, which varies across loci and individuals.

Here we introduce scPloidyR, an R package that implements a HMM that jointly models read depth and B-allele frequency at amplicon resolution for single-cell CNV inference from Tapestri data. We compare scPloidyR with the established karyotapR GMM method through systematic simulation studies that evaluate performance across varying levels of simulation parameters (BAF noise, variant density, amplicon density, sample size, and heterozygosity rate) that govern the information content available to each method and through application to a public Tapestri cell mixture dataset. Our results characterize the conditions under which joint depth-BAF modeling improves copy number calling and identify regimes where depth-only inference remains preferable.

## Results

### Simulation study 1

Simulation study 1 mixed multiple cell populations across copy-number states to approximate real datasets with heterogeneous karyotypes (see Methods). Across ten replicates, scPloidyR consistently outperformed karyotapR on all discrimination-focused metrics. Mean accuracy for scPloidyR was 0.858, compared with 0.467 for karyotapR and 0.788 for the naive diploid predictor (Figure 1A); however, naive accuracy reflects the dominance of diploid segments and does not indicate true CNV detection ability. scPloidyR achieved substantially higher macro-F1 (0.472 vs. 0.264; Figure 1B) and balanced accuracy (0.521 vs. 0.304; Figure 1C), demonstrating improved performance across all copy-number classes rather than only the dominant class. Importantly, scPloidyR maintained high alteration F1 (0.902 vs. 0.383 for karyotapR; Figure 1D), whereas the naive baseline had zero alteration F1. The ground truth, as expected, achieved perfect scores across all metrics.

**Figure 1:**
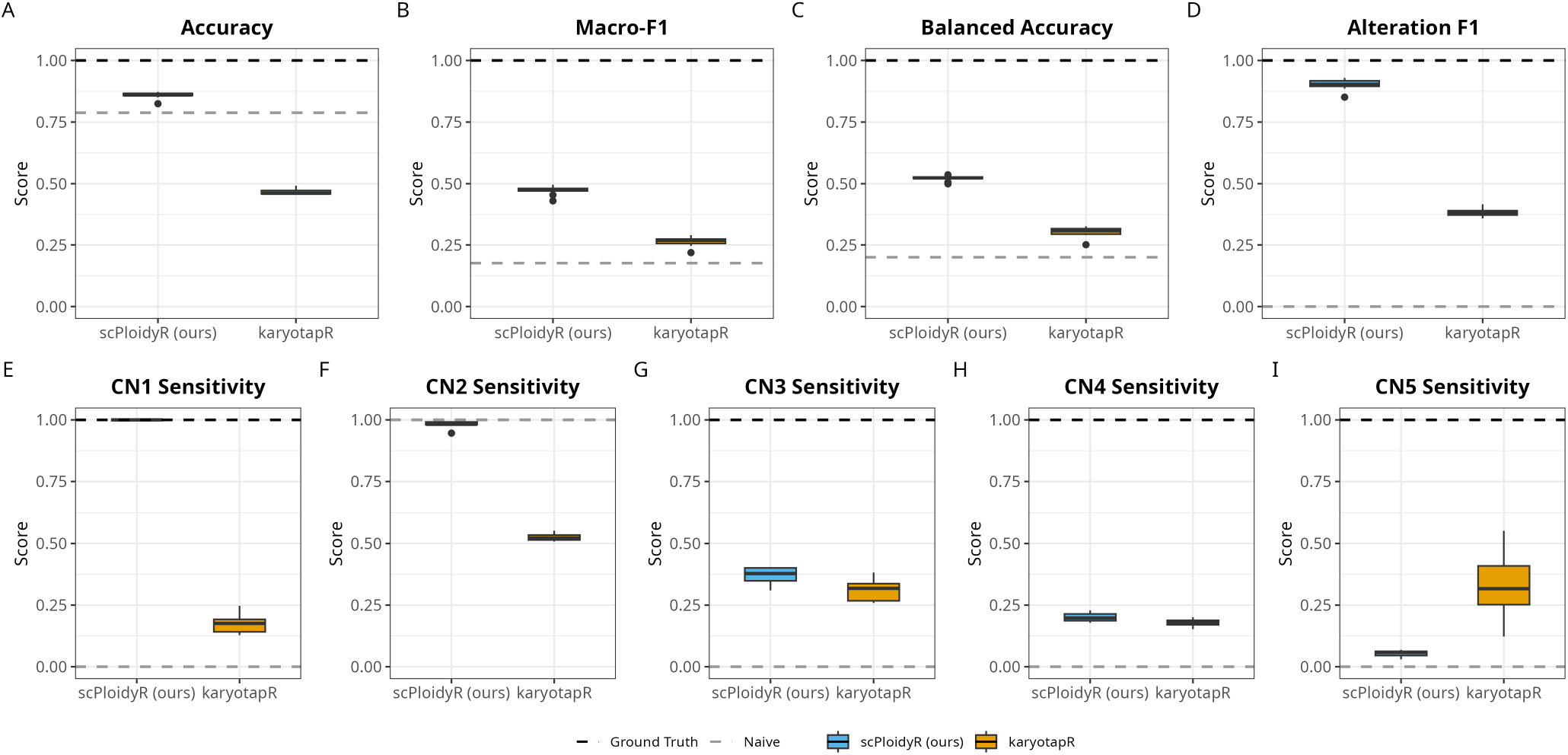
Simulation study 1: comparison of scPloidyR and karyotapR (GMM) across copy number states 1–5 over ten independent replicates, with dashed lines for the ground truth (upper bound) and naive diploid predictor (always CN = 2). (A) Accuracy, (B) macro-F1, (C) balanced accuracy, (D) alteration F1, (E–I) per-class sensitivity for CN states 1–5.

Per-class sensitivity analysis reveals how each method performs across individual copy number states. For singlecopy deletions (CN 1), scPloidyR achieved perfect sensitivity (1.000) compared with 0.175 for karyotapR (Figure 1E). For the diploid state (CN 2), scPloidyR achieved 0.982 sensitivity compared with 0.526 for karyotapR (Figure 1F), indicating that scPloidyR correctly identifies the majority class far more reliably. For single-copy gains (CN 3), scPloidyR modestly outperformed karyotapR (0.371 vs. 0.309; Figure 1G). At higher ploidy states, sensitivity dropped for both methods: CN 4 sensitivity was 0.200 for scPloidyR versus 0.180 for karyotapR (Figure 1H), while for CN 5, karyotapR showed higher sensitivity than scPloidyR (0.331 vs. 0.054; Figure 1I), suggesting that the GMM approach may better distinguish extreme gains. The naive diploid predictor, by construction, has zero sensitivity for all non-diploid states. These per-class results show that aggregate metrics such as accuracy can obscure important differences in method behavior across copy number states. Full numerical results for all metrics are provided in Supplementary File S5.

### Simulation study 2

Simulation study 2 varied five parameters one at a time, BAF noise, variant density, amplicon density, sample size, and heterozygosity rate, with three replicates per condition for a total of 108 simulation runs (see Methods). Each condition was evaluated separately for additions (CN 3) and deletions (CN 1). Across most conditions, scPloidyR outperformed karyotapR, but the magnitude of the gap varied substantially by parameter, and karyotapR achieved higher accuracy when BAF information was absent, i.e. zero variants or zero heterozygosity. The subsections below describe each tested variable in detail.

#### BAF noise

Increasing BAF noise strongly degraded scPloidyR performance while leaving karyotapR essentially unchanged. For additions, scPloidyR accuracy fell from 0.990 (sd = 6) to 0.945 (sd = 9) to 0.787 (sd = 12), with macro-F1 declining in parallel from 0.983 to 0.904 to 0.716 and alteration F1 from 0.983 to 0.895 to 0.606. karyotapR accuracy remained flat at approximately 0.66 across all noise levels (0.657, 0.662, 0.665), and its alteration F1 remained near 0.37–0.39. A sharp inflection occurred between sd = 9 and sd = 12, where scPloidyR’s balanced accuracy dropped from 0.870 to 0.717. For deletions, the pattern was similar but less severe: scPloidyR accuracy declined from 1.000 to 0.984 to 0.852, while karyotapR remained near 0.69–0.71. These results indicate that scPloidyR’s advantage depends heavily on BAF signal quality, with high noise levels reducing its ability to discriminate copy-number states (Figure 2).

**Figure 2:**
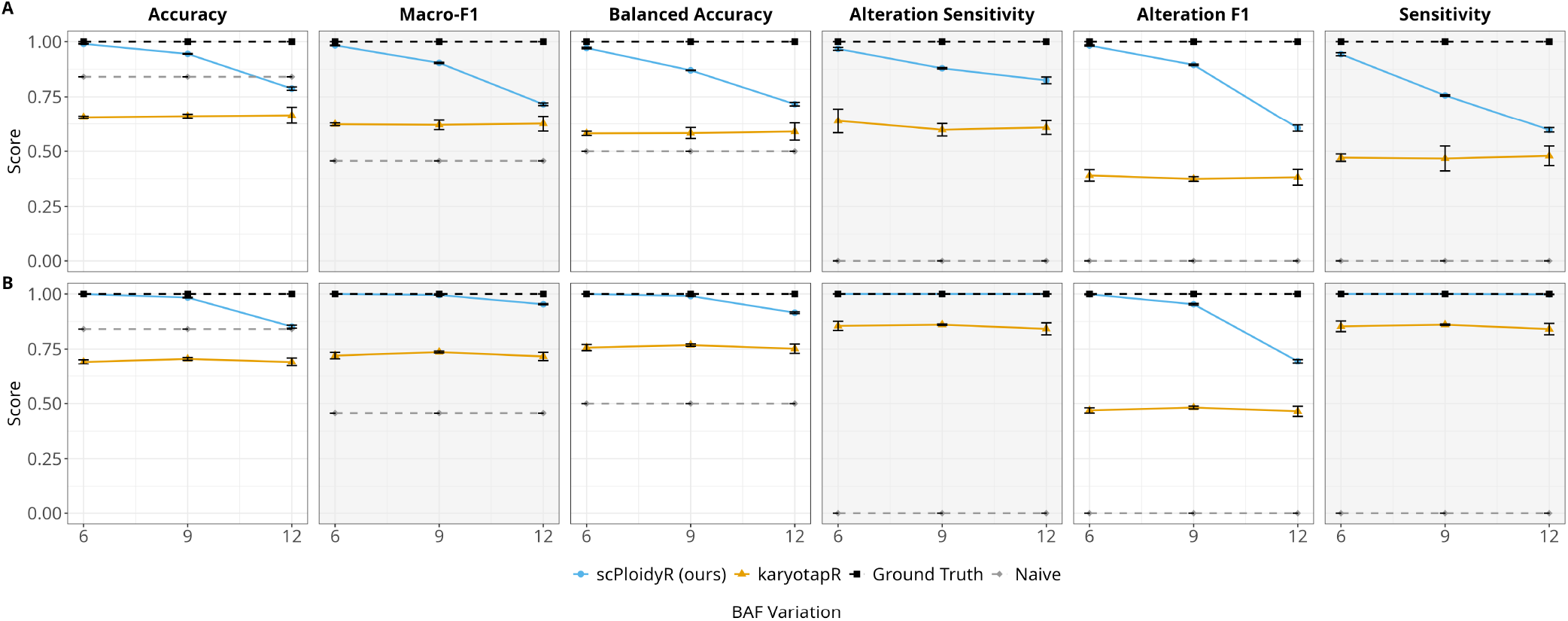
Simulation study 2: BAF noise univariate analysis. (A) Copy number gains (CN = 3); (B) copy number losses (CN = 1). Each panel plots a performance metric as a function of BAF noise (standard deviation of allelefrequency perturbation) with three replicates per condition. Lines represent scPloidyR (blue), karyotapR (GMM, orange), the ground truth (black), and the naive diploid predictor (grey).

#### Variant density

Variant density, heterozygous SNPs per amplicon, produced the largest change in scPloidyR performance. For additions, scPloidyR accuracy jumped from 0.548 with zero variants to 0.899 with one variant per amplicon. Performance continued to improve with additional variants (0.941 at 3, 0.963 at 5) but with diminishing returns. Alteration F1 showed the same pattern: 0.401 (0 variants), 0.794 (1 variant), 0.888 (3 variants), 0.945 (5 variants). karyotapR was unaffected by variant count, with accuracy between 0.656 and 0.681 across all levels. With zero variants, i.e. depthonly mode, scPloidyR lost its BAF advantage entirely and performed below the naive baseline in accuracy (0.548 vs. 0.841). For deletions, the zero-variant deficit was smaller but karyotapR again outperformed scPloidyR (0.720 vs. 0.668), and adding even one variant restored near-perfect scPloidyR performance (0.961) (Figure 3).

**Figure 3:**
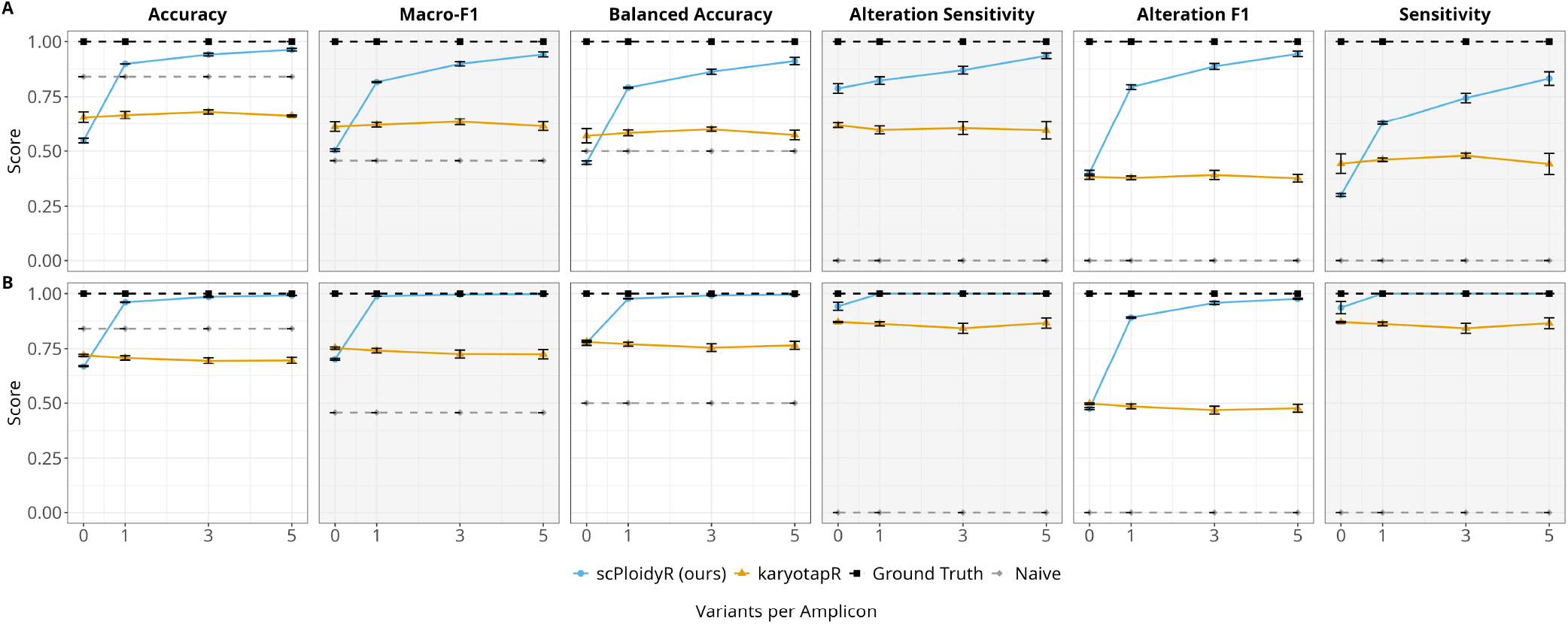
Simulation study 2: variant density univariate analysis. (A) Copy number gains (CN = 3); (B) copy number losses (CN = 1). Performance metrics plotted as a function of heterozygous SNPs per amplicon (0, 1, 3, 5). Layout and conventions as in Figure 2.

#### Amplicon density

Both methods improved with more amplicons per chromosome, but the pattern differed. karyotapR showed the larger relative gain for additions, with accuracy rising from 0.495 (2 amplicons) to 0.661 (4 amplicons) to 0.728 (6 amplicons), a 47% relative improvement. scPloidyR improved more modestly in absolute terms (0.871 to 0.950 to 0.967) since it was already performing well at low amplicon counts. Alteration F1 for scPloidyR rose from 0.772 to 0.908 to 0.949, while karyotapR improved from 0.305 to 0.377 to 0.445. The improvement was monotonic for both methods with no inflection point, suggesting that additional spatial resolution continues to benefit copy-number inference. For deletions, scPloidyR was already near-perfect at 2 amplicons (accuracy 0.932) and reached 0.994 at 6 amplicons (Figure 4).

**Figure 4:**
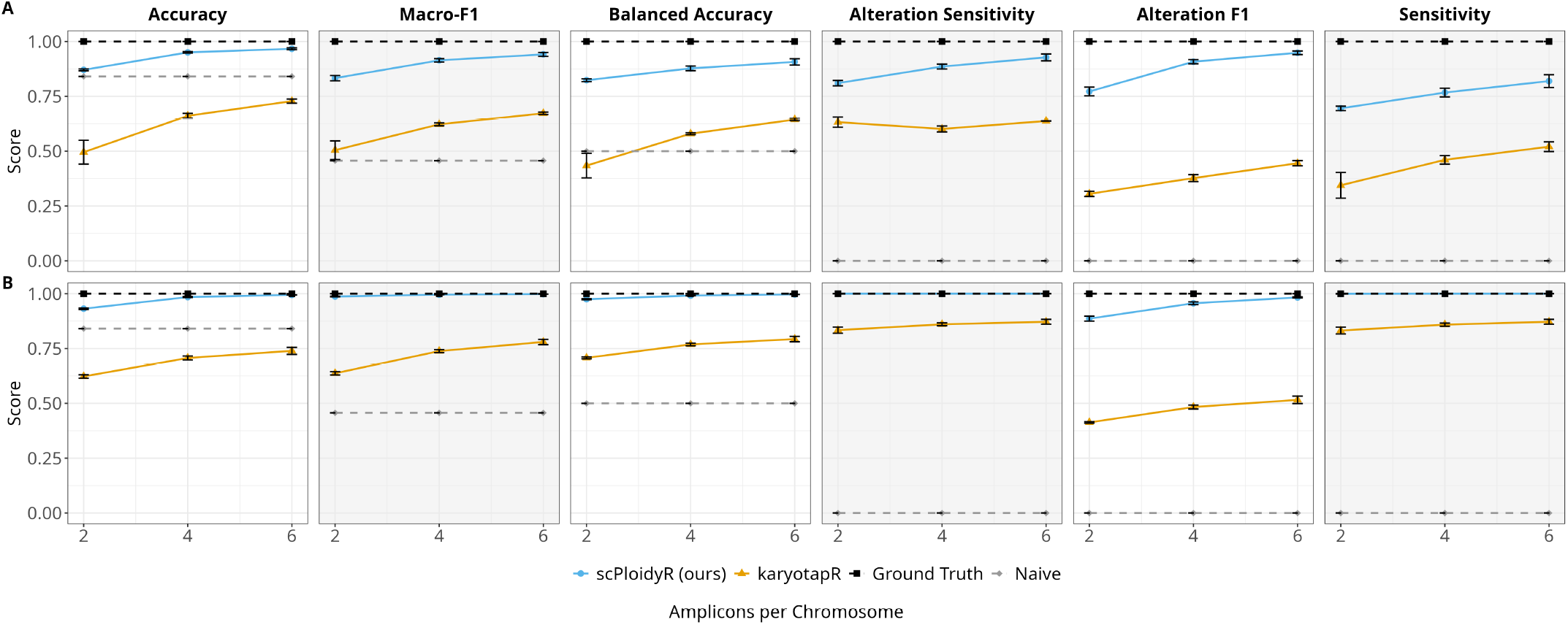
Simulation study 2: amplicon density univariate analysis. (A) Copy number gains (CN = 3); (B) copy number losses (CN = 1). Performance metrics plotted as a function of amplicons per chromosome (2, 4, 6). Layout and conventions as in Figure 2.

#### Sample size

Sample size, cells per group, had minimal impact on either method. For additions, scPloidyR accuracy was 0.945 at 50 cells, 0.948 at 100 cells, and 0.942 at 200 cells; macro-F1 varied by less than 0.012 across conditions (0.900–0.912). karyotapR showed similarly flat performance (accuracy 0.655–0.669). For deletions, the same stability held: scPloidyR accuracy ranged from 0.983 to 0.987 and karyotapR from 0.691 to 0.709. These results indicate that reference panel estimation is stable across different number of cells within a group (Figure 5).

**Figure 5:**
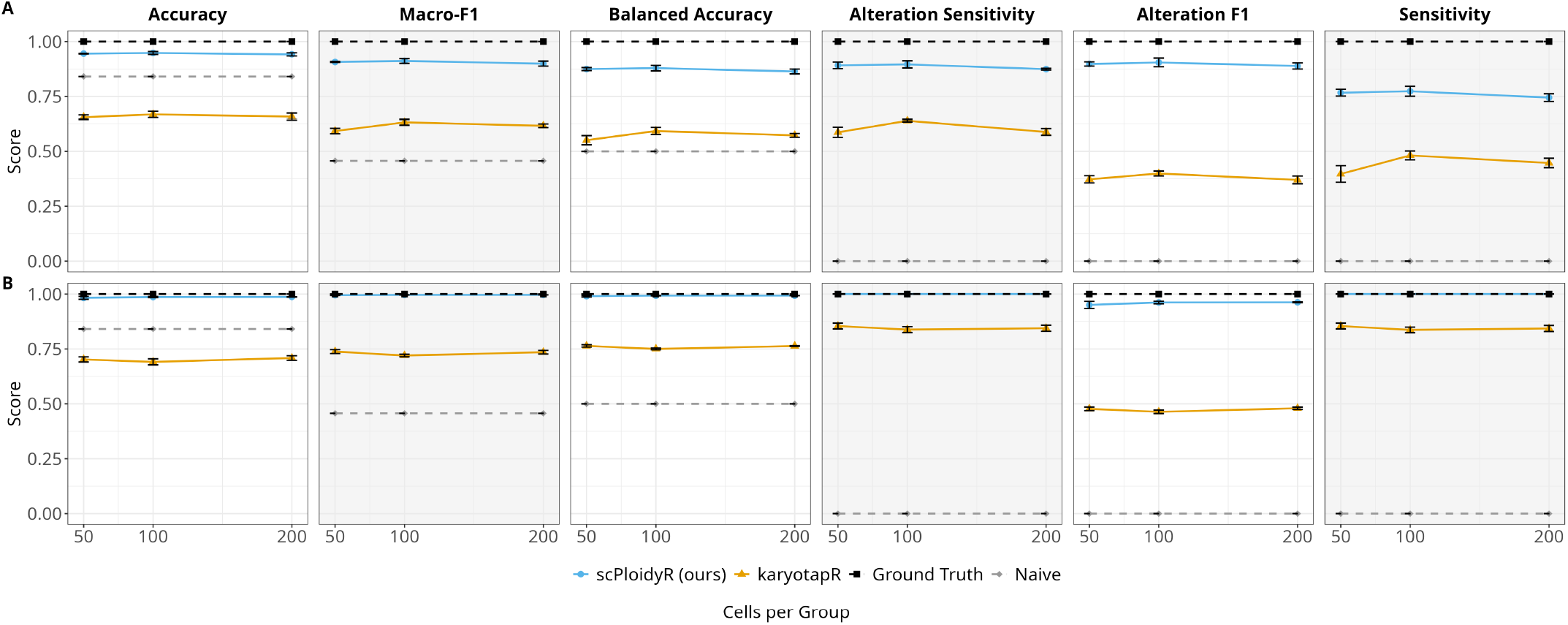
Simulation study 2: sample size univariate analysis. (A) Copy number gains (CN = 3); (B) copy number losses (CN = 1). Performance metrics plotted as a function of cells per group (50, 100, 200). Layout and conventions as in Figure 2.

#### Heterozygosity rate

Heterozygosity rate produced the largest performance swing for scPloidyR. For additions, accuracy rose from 0.569 at 0% heterozygosity to 0.655 (25%), 0.772 (50%), 0.843 (75%), and 0.949 (100%). Alteration F1 showed a similarly steep trajectory: 0.395, 0.467, 0.584, 0.681, and 0.905 across the same levels. The curve was non-linear, with steeper gains at higher heterozygosity rates. karyotapR remained stable across all heterozygosity levels (accuracy 0.660–0.682, alteration F1 0.371–0.389). At 0% heterozygosity, karyotapR outperformed scPloidyR (0.662 vs. 0.569), demonstrating that depth-only inference surpasses the HMM when no allelic information is available. At 25% heterozygosity, karyotapR still narrowly outperformed scPloidyR (0.661 vs. 0.655). For deletions, heterozygosity had a qualitatively similar but less extreme effect: scPloidyR accuracy rose from 0.630 (0%) to 0.988 (100%). At 0% heterozygosity for deletions, karyotapR again outperformed scPloidyR (0.695 vs. 0.630) (Figure 6).

**Figure 6:**
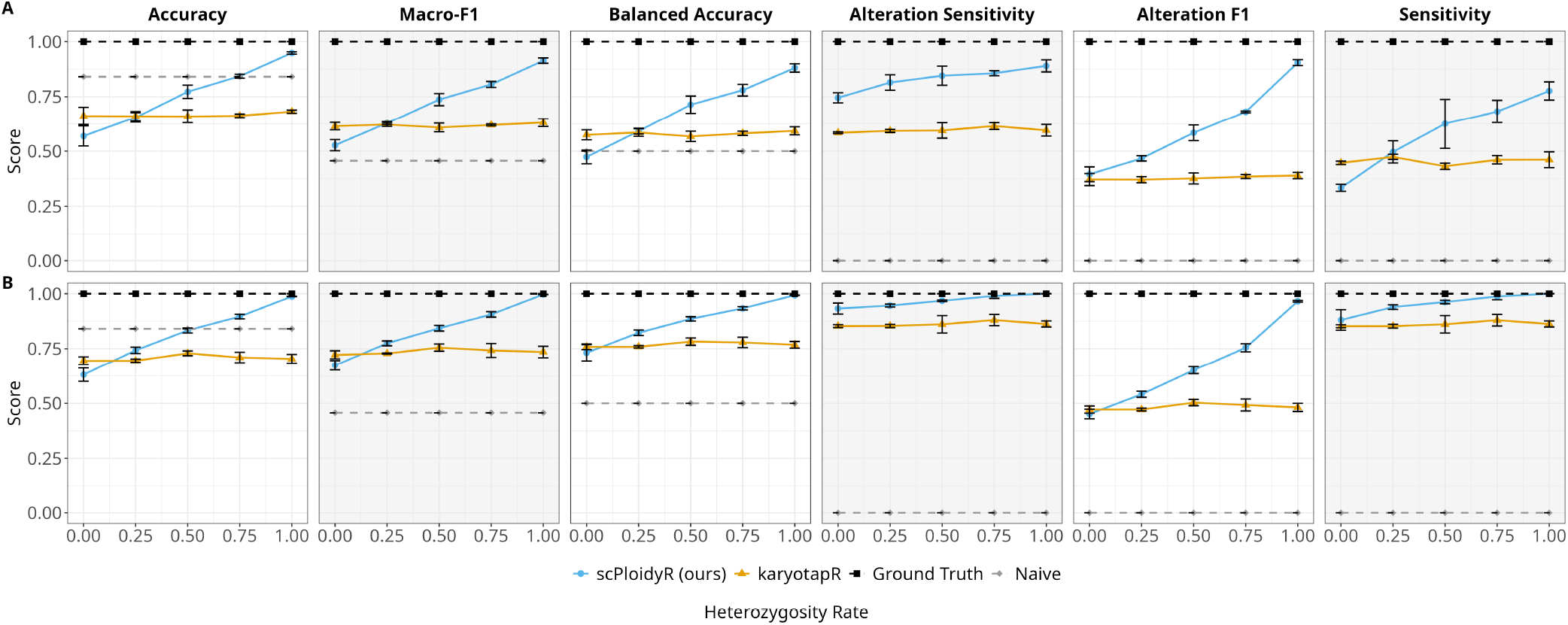
Simulation study 2: heterozygosity rate univariate analysis. (A) Copy number gains (CN = 3); (B) copy number losses (CN = 1). Performance metrics plotted as a function of heterozygosity rate (0%, 25%, 50%, 75%, 100%). Layout and conventions as in Figure 2.

#### Summary of simulation study 2 trends

Across the five tested variables, scPloidyR performance was primarily driven by BAF-related parameters, namely heterozygosity rate, variant density, and BAF noise, which jointly determine the information content of allelefrequency signals. When BAF information was absent, i.e. zero variants per amplicon or 0% heterozygosity, karyotapR outperformed scPloidyR for both additions and deletions. karyotapR was mainly sensitive to amplicon density, which governs spatial resolution for depth-based inference. Sample size had minimal impact on either method. Deletions were consistently easier to detect than additions for both methods, likely because monosomy produces a larger departure from the diploid baseline than trisomy. Detailed metric values for all conditions are provided in Supplementary Files S6 (gains) and S7 (losses).

### Application to real data

We applied both methods to the public Tapestri cell mixture dataset (see Methods). In the absence of ground truth, the comparison focuses on concordance and biological plausibility. The karyotapR (GMM) and scPloidyR copynumber heatmaps (Figure 7A and 7B) provide direct qualitative comparisons across the five cell-line populations.

**Figure 7:**
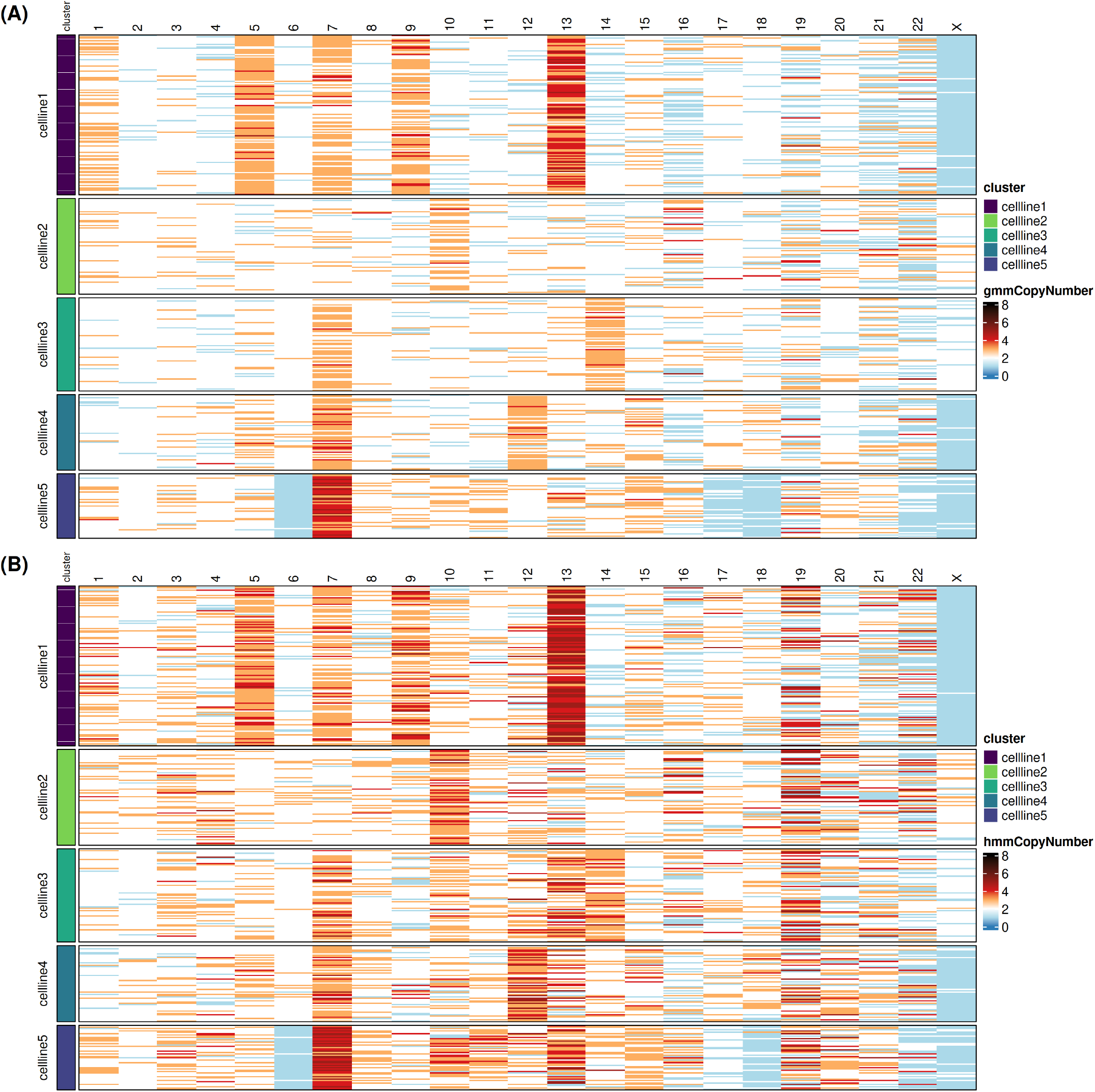
Single-cell copy number heatmaps from the Tapestri five-cell-line mixture dataset. (A) karyotapR GMM copy number calls. (B) scPloidyR HMM copy number calls. Rows represent individual cells grouped by cell line identity (color bar, left); columns represent chromosomes 1–22 and X. Copy number states are encoded by a diverging color scale (blue: loss, white: diploid, orange/red: gain). Cell line 2 (RPE1) served as the near-diploid reference with known trisomy 10q.

Both methods recover concordant large-scale karyotypic structures across all five cell lines. Cell line 1 displays extensive gains on chromosomes 1, 5, 7, 9, and 13; the RPE1 reference (cell line 2) is predominantly diploid apart from the expected trisomy 10 (Mays et al., 2025); and cell lines 3–5 each exhibit distinct but consistent alteration patterns in both heatmaps. The spatial distribution of gains (orange/red) and losses (blue) is broadly similar, confirming that both approaches detect the major CNV events in this dataset.

Despite this overall concordance, several qualitative differences are apparent. Visually, chromosome 19 stands out as one of the most clearly divergent regions between the two methods. karyotapR produces more uniformly diploid (white) calls on chromosome 19 in all cell lines, whereas scPloidyR shows more scattered gain calls in the same region, suggesting a more flexible copy number call in the HMM based model using BAF signals. Second, for the X chromosome in cell lines 1, karyotapR assigns noticeably more heterogeneous copy number states across cells, while scPloidyR yields more uniform calls within this cell population. The difference between such region may indicate a consistant copy number call driven by their BAF signals.

### Summary of comparative performance

Across both simulation study 1 and simulation study 2, scPloidyR generally improves class-balanced performance and alteration detection relative to karyotapR, while the naive diploid predictor achieves misleadingly elevated accuracy but fails to detect CNVs. However, when BAF information is unavailable, either due to zero heterozygous variants per amplicon or 0% heterozygosity, karyotapR outperforms scPloidyR. Overall, the results indicate that joint modeling of depth and BAF provides a clear advantage for single-cell CNV calling in heterogeneous populations, but that depth-only methods remain preferable when allelic information is absent.

## Discussion

This study demonstrates that jointly modeling read depth and B-allele frequency within an HMM framework substantially improves single-cell copy number calling from targeted DNA sequencing panels. Across both simulation studies, scPloidyR outperformed the depth-only karyotapR GMM on macro-F1, balanced accuracy, and alteration F1, and application to a real Tapestri cell mixture dataset confirmed that the joint model produces more spatially coherent copy number profiles.

The primary advantage of scPloidyR lies in exploiting allelic information invisible to depth-only methods. BAF distinguishes copy number states that produce similar total read counts, such as different trisomy genotypes that share the same depth but yield distinct BAF values (Wang et al., 2007; Zaccaria & Raphael, 2021). The variant density experiment directly demonstrates this: adding just one heterozygous variant per amplicon increased scPloidyR accuracy from 0.548 to 0.899 for gains, confirming that even sparse allelic information provides substantial discriminative power. The per-chromosome HMM structure further contributes by enforcing spatial coherence, smoothing isolated noisy observations toward the local consensus state and reducing false positive calls in regions without true alterations.

However, scPloidyR is heavily dependent on BAF availability and quality. When heterozygous variants are absent, scPloidyR loses its allelic signal and underperforms karyotapR. The BAF noise experiment revealed a second vulnerability: as noise increased from sd = 9 to sd = 12, scPloidyR accuracy for gains dropped from 0.945 to 0.787, while karyotapR remained stable. These results indicate that the quality, not merely the presence, of allelic information determines whether joint modeling improves upon depth-only inference. As a practical guideline, scPloidyR should be preferred when at least one heterozygous variant per amplicon is available and BAF noise is moderate; karyotapR is the safer choice when allelic information is sparse or unreliable.

Several limitations should be noted. Both methods showed reduced sensitivity for high copy number states (CN 4 and CN 5), suggesting that neither reliably resolves high-ploidy states. The simulation framework employed simplified ground truth designs with uniform copy number states; real tumors exhibit more complex patterns including subclonal alterations and focal events (Navin et al., 2011; Zack et al., 2013) that may affect performance differently. Additionally, the application study lacked ground truth copy number profiles, limiting the comparison to qualitative concordance rather than quantitative accuracy.

Despite these limitations, our results establish that joint depth-BAF modeling via scPloidyR provides a clear and substantial improvement over depth-only approaches for single-cell CNV calling when allelic information is available. By combining the discriminative power of BAF with the spatial smoothing of a per-chromosome HMM, scPloidyR offers a principled and effective framework for copy number inference from targeted single-cell DNA sequencing panels.

## Methods

### Data normalization and reference specification

Normalized read counts are computed using Mission Bio’s mb scheme. Let *d*_*i,c*_ denote the raw read count for amplicon *i* in cell *c* (*c* = 1, …, *C*, where *C* is the total number of cells), and *A* the number of amplicons. First, “good barcodes” are defined as cells with total counts exceeding 10% of *S*_(11)_, the 11th-largest total count across all cells:

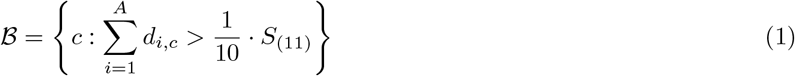

Second, counts are adjusted for cell depth by dividing by the cell mean (plus a stabilizing constant):

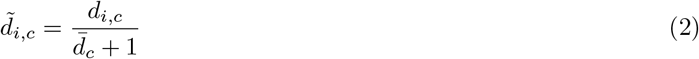

where 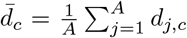. Third, amplicon-level normalization uses medians across good barcodes and scales to the diploid baseline:

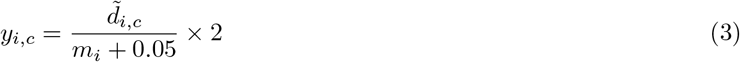

where 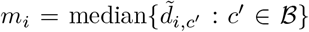. Reference cells are cell clusters defined by users with known ploidy (typically diploid) that define the baseline copy number profile. karyotapR accepts a `control.copy.number` template; sc-PloidyR uses `reference_cn`. Non-diploid regions in reference cells (for example, trisomy) are explicitly encoded in these templates.

### Copy number inference models

**karyotapR (GMM)**. For each probe *i*, a Weibull distribution is fitted to normalized counts in reference cells. The Weibull distribution was selected because it provides a good fit for the observed distribution of normalized amplicon counts (Mays et al., 2025). Synthetic cells are generated for each copy number state *k* by scaling the Weibull scale parameter:

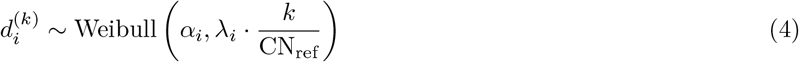

with shape *α*_*i*_ and scale *λ*_*i*_ estimated per probe and CN_ref_ typically 2. Because each draw from Eq. 4 represents one synthetic cell rather than an observed cell, the cell index *c* is implicit. Probe-level values are summarized to segments (chromosome arms or whole chromosomes) using the median. Gaussian components 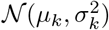 are fitted to each copy number class, and real cells are classified by posterior probability under equal priors:

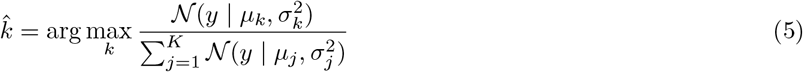

where *y* is the segment-level copy number value.

**scPloidyR**. scPloidyR models the ordered amplicons along each chromosome as emissions from a Markov chain with hidden copy number states **z** = (*z*_1_, …, *z*_*n*_), where *z*_*i*_ ∈ {1, 2, …, *K*} with *K* = 5. Transitions are governed by a persistence matrix with rare breakpoints:

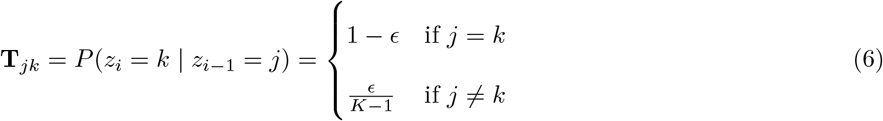

with *ϵ* initialized at 10^−3^. The emission probability factorizes into depth and BAF likelihoods, where 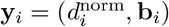 is the observation vector at amplicon *i*, consisting of the normalized read depth 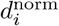 and the vector of minor allele frequencies **b**_*i*_:

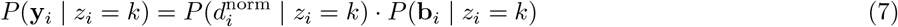

Depth is modeled as Gaussian on the normalized scale with state-dependent means:

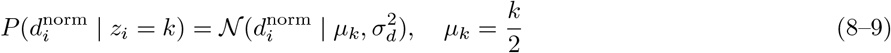

where 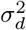 is the variance of normalized depth across amplicons (subscript *d* for depth). Input depths 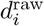 (MB-normalized counts from Eq. 3) are normalized by per-amplicon baselines learned from reference cells:

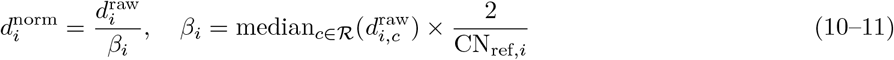

where *ℛ* are reference cells and CN_ref,*i*_ permits non-diploid reference regions.

For allele frequency, each amplicon *i* provides *V*_*i*_ variants. Raw B-allele frequencies are converted to minor allele frequencies (MAF) via *b* = min(BAF, 1 − BAF), yielding observed values **b**_*i*_ = (*b*_*i*,1_, …, *b*_*i,V*_*i*) on [0, 0.5]. The likelihood marginalizes over genotypes compatible with copy number state *k*, where *𝒢*_*k*_ is the set of possible genotypes for copy number state *k*:

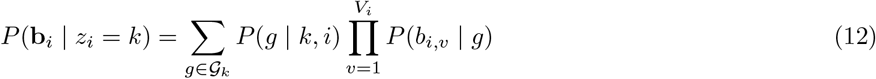

Genotypes are all combinations of *k* alleles (A or B). The expected minor allele frequency (MAF) for a genotype with *m* minor-allele copies is min(*m, k* − *m*)*/k*, yielding representative values of 0 (homozygous), 1*/*3 for *k* = 3 heterozygotes, 1*/*4 and 1*/*2 for *k* = 4 heterozygotes, and analogous values for general *k*. Genotype priors are anchored by the per-amplicon heterozygosity rate *h*_*i*_ estimated from reference cells; for diploid loci:

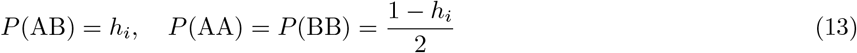

with higher copy number priors interpolated between homozygous and heterozygous templates. Each MAF observation is modeled by a truncated normal on [0, 0.5]:

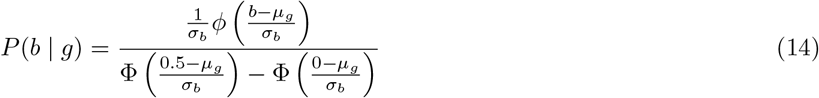

where *µ*_*g*_ is the expected MAF for genotype *g*.

#### Reference parameter learning and EM

scPloidyR learns per-amplicon depth baselines *β*_*i*_, depth variance 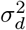, BAF standard deviation *σ*_*b*_, and heterozygosity rates *h*_*i*_ from reference cells. Model parameters 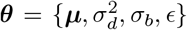 are then estimated using Baum–Welch with forward–backward posteriors

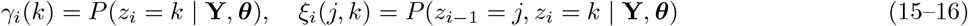

where **Y** denotes the complete set of observations across all amplicons, and standard M-step updates for *µ*_*k*_, 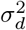, *σ*_*b*_, and *ϵ*:

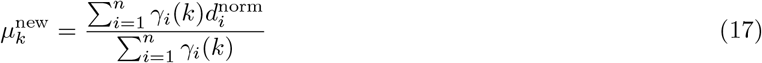

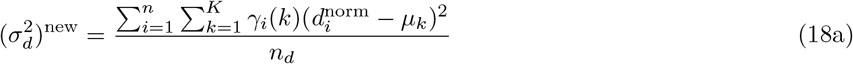

where *n*_*d*_ is the number of amplicons with observed depth, and

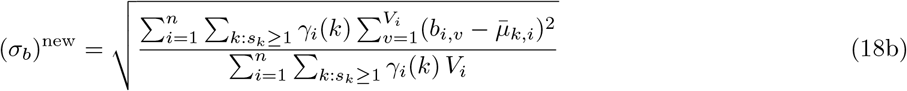

where 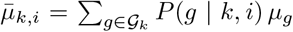 is the prior-weighted expected MAF for state *k* at amplicon *i*, and the sums are restricted to informative states (*s*_*k*_ ≥ 1) with observed variants (*V*_*i*_ *>* 0).

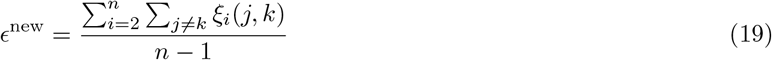

When heterozygosity is low (mean heterozygosity 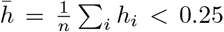), depth means can collapse. We therefore regularize toward prior means 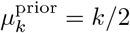 using a weight *w* that increases as *h* decreases:

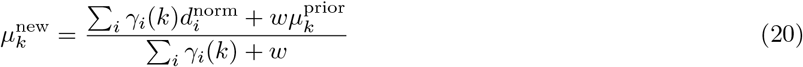

EM iterates until parameter change is *<* 10^−5^ or 50 iterations. Final copy number paths are obtained by Viterbi decoding:

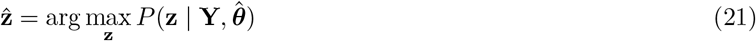

To enforce chromosome boundaries, scPloidyR fits independent chains per chromosome and concatenates per-chromosome Viterbi paths. Amplicon-level calls are aggregated to segments using the mode (scPloidyR) or median (karyotapR):

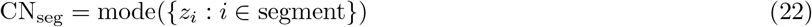

### Simulation study

We evaluated methods on synthetic data with known ground truth (i.e., known copy-number assignments) in two simulation studies. **Simulation study 1** used a balanced multi-state design with six cell populations of equal size: copy number (CN) = 1 (monosomy), CN=2 (disomy/diploid, reference), CN=2 (disomy, mixed genotypes), CN=3 (trisomy), CN=4 (tetrasomy), and CN=5 (pentasomy). Fifty percent of chromosomes were randomly altered in nonreference groups across 22 chromosomes with 4 amplicons per chromosome, 3 variants per amplicon, BAF noise (the standard deviation of Gaussian noise added to simulated allele frequencies, denoted sd) of 9, and 100% heterozygosity. Ten independent replicates were run with parameter jitter (cells ± 10%, BAF noise ± 15%). **Simulation study 2** used a multi-cell line design with two populations — a diploid reference and one altered cell line — with 30% of chromosomes altered. Five parameters were varied one at a time (BAF noise, variants per amplicon, amplicons per chromosome, cells per group, and heterozygosity rate), each tested for both gains (CN=3) and losses (CN=1), with three independent replicates per condition (108 total runs). Baseline values were BAF sd=9, 3 variants per amplicon, 4 amplicons per chromosome, 100 cells per group, and 100% heterozygosity. Table 1 summarizes the ground truth design for both simulation studies.

**Table 1a.** Simulation study 1: synthetic cell populations. Ten replicates with parameter jitter (cells ± 10%, BAF noise ± 15%).

| Population | CN | Cells | Chr. Altered | Replicates |
| --- | --- | --- | --- | --- |
| CN=1 | 1 | 80 | 11/22 | 10 |
| CN=2 (reference) | 2 | 80 | 0/22 | 10 |
| CN=2 (mixed) | 2 | 80 | 0/22 | 10 |
| CN=3 | 3 | 80 | 11/22 | 10 |
| CN=4 | 4 | 80 | 11/22 | 10 |
| CN=5 | 5 | 80 | 11/22 | 10 |

CN=2 (reference) cells are diploid across all chromosomes and serve as the reference for normalization. CN=2 (mixed) cells are also diploid but carry heterogeneous genotype configurations to test genotype-driven BAF variation without copy number changes.

Study 2 varied parameters one at a time: BAF sd (6, 9, 12), variants/amplicon (0, 1, 3, 5), amplicons/chr (2, 4, 6), cells/group (50, 100, 200), heterozygosity rate (0%, 25%, 50%, 75%, 100%). Baseline values: 4 amplicons/chr, 3 variants/amplicon, BAF sd=9, 100 cells/group, 100% heterozygosity. Tables 1b and 1c show baseline values; each varied parameter replaced one baseline value at a time (36 combinations, 3 replicates each, 108 total runs). A complete summary of simulation parameters and conditions is provided in Supplementary File S1.

**Table 1b.** Simulation study 2 (copy number gain): synthetic cell populations.

| Population | CN | Cells | Chr. Altered | Replicates |
| --- | --- | --- | --- | --- |
| Diploid reference | 2 | 100 | 0/22 | 3 |
| Altered (gain) | 3 | 100 | 7/22 | 3 |

**Table 1c.** Simulation study 2 (copy number loss): synthetic cell populations.

| Population | CN | Cells | Chr. Altered | Replicates |
| --- | --- | --- | --- | --- |
| Diploid reference | 2 | 100 | 0/22 | 3 |
| Altered (loss) | 1 | 100 | 7/22 | 3 |

#### Read count simulation

For amplicon *i* and cell *c*, counts were sampled from a Weibull distribution with mean proportional to copy number:

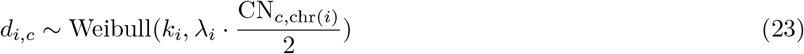

with *k*_*i*_ ~ |*𝒩* (2, 0.8^2^)| + 0.3. The scale parameter *λ*_*i*_ was constructed to be inversely related to *k*_*i*_ and normalized to [0.5, 2.5], reproducing heterogeneous amplicon capture efficiencies observed in Tapestri data. Parameters (*k*_*i*_, *λ*_*i*_) were shared across cell lines to preserve realistic amplicon-specific biases.

#### BAF simulation

For each variant, a genotype was assigned according to the chromosome genotype and heterozygosity rate, the expected MAF was computed, and Gaussian noise with standard deviation equal to the BAF noise parameter (sd) was added, with truncation to [0, 1]. The number of variants per amplicon was sampled from user-defined options (with a default of 3 variants per amplicon in the baseline condition).

#### Univariate analyses

Simulation study 2 experiments used varied BAF noise (standard deviation 6, 9, 12), variants per amplicon (0, 1, 3, 5), amplicons per chromosome (2, 4, 6), cells per group (50, 100, 200), and heterozygosity rate (0%, 25%, 50%, 75%, 100%). Each combination was tested for both gains (CN=3) and losses (CN=1), with three independent replicates.

#### Evaluation metrics

Performance was assessed using accuracy

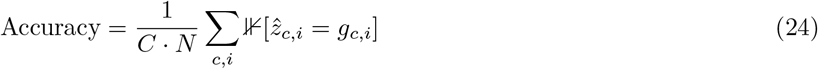

where *N* is the number of segments and ⊮[·] denotes the indicator function, macro-F1

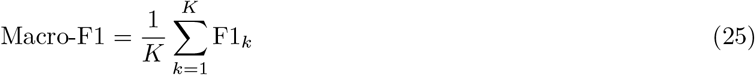

balanced accuracy

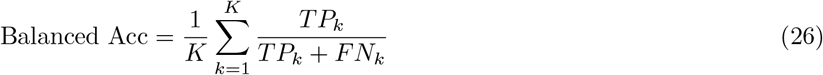

and per-class sensitivity

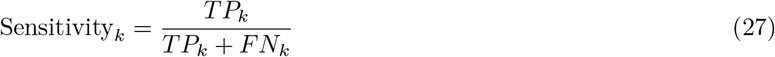

We additionally evaluated binary detection of any CNV (CN ≠ 2), using a naive diploid predictor (always CN=2) as a performance floor. Letting TP and FN denote true positives and false negatives under the binary classification where “positive” is any alteration (CN ≠ 2) and “negative” is diploid (CN = 2). Because this is a binary classification (altered vs. diploid), class-specific subscripts are not needed. Alteration sensitivity is

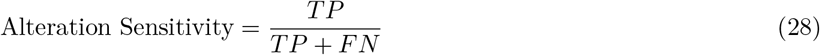

and alteration F1 combines sensitivity with alteration precision Precision_alt_ = *TP/*(*TP* + *FP*):

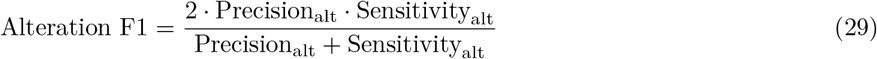

Each metric captures a distinct facet of copy number calling performance. Overall accuracy provides a straightforward measure of correctness but can be misleading when diploid segments dominate the profile, inflating scores for methods that default to CN=2. Macro-F1 addresses this class imbalance by averaging F1 scores equally across all CN states, ensuring that rare but clinically important states such as deletions and high-level gains contribute as much as the diploid majority. Balanced accuracy shares this motivation, averaging per-class recall so that majority-class dominance does not inflate aggregate scores. Per-class sensitivity reveals which specific CN states a method can and cannot detect, exposing failure modes that summary statistics may obscure. Alteration sensitivity measures the ability to detect any non-diploid event, the core clinical task of CNV screening, while alteration F1 balances detection of true alterations against false positives—relevant for clinical decision-making where both missed calls and spurious calls carry consequences.

#### Confusion matrices

For each simulation replicate and method, we computed a multi-class confusion matrix **M** ∈ Z^*K*×*K*^ over copy number states {1, …, *K*}. Predicted and ground truth copy number matrices were flattened to vectors; each element was tabulated into entry *M*_*jk*_, the count of observations with true state *j* and predicted state *k*. Per-class precision, recall (sensitivity), and F1 scores were derived from the confusion matrix as Precision_*k*_ = *M*_*kk*_*/∑* _*j*_ *M*_*jk*_, Recall_*k*_ = *M*_*kk*_*/∑* _*j*_ *M*_*kj*_, and F1_*k*_ = 2 · Precision_*k*_ · Recall_*k*_*/*(Precision_*k*_ + Recall_*k*_), respectively. Full confusion matrices for all simulation conditions are provided in Supplementary Files S2–S4.

### Application study

We applied both methods to the public karyotapR tutorial dataset (Mays et al., 2025; Mays & Davoli, 2023), a mixture of five human cell lines sequenced with the Mission Bio Tapestri CO261 panel (317 amplicons across chromosomes 1–19). Cell populations were identified by allele-frequency-based clustering using PCA, UMAP, and DBSCAN. Cell line 2 (RPE1) served as the near-diploid reference; its known trisomy of chromosome 10q was encoded as `reference_cn` = c(“10” = 3) in both methods.

Preprocessing removed amplicons with near-zero median counts in reference cells (9–11 probes, analysis-dependent), normalized counts with the mb scheme, and extracted BAF values from the `alleleFrequency` assay with variants mapped to amplicons by genomic coordinates. karyotapR inferred continuous copy number values per amplicon, smoothed to segments, and fit a GMM with components 1–5. scPloidyR learned reference parameters, fit a perchromosome HMM with states 1–5, and aggregated amplicon calls by segment mode. Without ground truth, we evaluated concordance between methods, consistency with known karyotypes.

### Software application

All analyses were implemented in R and are available in Github. karyotapR (version 1.0.1) was used as distributed through CRAN. The scPloidyR package is available at https://github.com/dpei/scPloidyR and the simulation study code is available at https://github.com/dpei/scPloidyR_simulation.

## Acknowledgements

The author(s) declare that financial support was received for the research and/or publication of this article. Research reported was primarily supported by the National Institute of Diabetes and Digestive and Kidney Diseases award R01 DK132320, Kansas Center for Metabolism and Obesity Research (KC-MORE) (Supported by the National Institute of General Medical Science award P20 GM144269), the National Cancer Institute (NCI) Cancer Center Support Grant P30 CA168524, the Kansas Institute for Precision Medicine COBRE (Supported by the National Institute of General Medical Science award P20 GM130423), the Kansas IDEA Network Of Biomedical Research Excellence (P20 GM103418), and the Department of Defense (HT94252410063).

## Notes

### Competing Interest Statement

The authors have declared no competing interest.

